# Alterations of the pro-survival Bcl-2 protein interactome in breast cancer at the transcriptional, mutational and structural level

**DOI:** 10.1101/695379

**Authors:** Simon Mathis Kønig, Vendela Rissler, Thilde Terkelsen, Matteo Lambrughi, Elena Papaleo

## Abstract

Apoptosis is an essential defensive mechanism against tumorigenesis. Proteins of the B-cell lymphoma-2 (Bcl-2) family regulates programmed cell death by the mitochondrial apoptosis pathway. In response to intracellular stresses, the apoptotic balance is governed by interactions of three distinct subgroups of proteins; the activator/sensitizer BH3 (Bcl-2 homology 3)-only proteins, the pro-survival, and the pro-apoptotic executioner proteins. Changes in expression levels, stability, and functional impairment of pro-survival proteins can lead to an imbalance in tissue homeostasis. Their overexpression or hyperactivation can result in oncogenic effects. Pro-survival Bcl-2 family members carry out their function by binding the BH3 short linear motif of pro-apoptotic proteins in a modular way, creating a complex network of protein-protein interactions. Their dysfunction enables cancer cells to evade cell death. The critical role in homeostasis and tumorigenesis coupled with progress in their structural elucidation, has led to consider pro-survival Bcl-2 proteins as therapeutic targets.

A better understanding of the transcriptomic level, mutational status and molecular mechanism underlying pro-survival Bcl-2 proteins in different cancer types, could help to clarify their role in cancer development and may guide advancement in drug discovery, targeting these proteins. Here, we shed light on pro-survival Bcl-2 proteins in breast cancer by proposing a ‘multiscale’ bioinformatic approach. We analyzed the changes in expression of the Bcl-2 proteins and their BH3-containing interactors, in breast cancer samples. We then studied, at the structural level, a selection of interactions, also accounting for effects induced by mutations found in the breast cancer samples. We identified the complexes between the up-regulated BCL2A1 and two down-regulated BH3-only candidates (HRK and NR4A1) as targets associated with reduced apoptosis in breast cancer samples, which could deserve future experimental validation. We predicted as damaging mutations altering protein stability L99R, M75R, along with Y120C as a possible allosteric mutation from an exposed surface to the BH3-binding site.

## Introduction

Apoptosis is a vital physiological process for embryogenesis, maintaining tissue homeostasis, discharging damaged, or infectious cells. Failures in apoptosis pathways may lead to carcinogenesis by favoring cell proliferation over cell death [1,2].

Apoptosis progresses through two discrete pathways: (i) intrinsic apoptosis (also called mitochondrial or stress-induced apoptosis), triggered by intracellular stresses, including oncogenic stress and chemotherapeutic agents [3], and (ii) extrinsic apoptosis, triggered by external stimuli detected by “death receptors” [4]. The intrinsic apoptotic pathway is governed by protein members of the B-cell lymphoma-2 (BCL-2) family, dictating the cellular decision making between cell survival or programmed cell death [5]. As a response to cellular stress, these proteins preserve the integrity of the cell or commits the cell to apoptosis by permeabilization of the outer mitochondrial membrane (OMM) and release of proteins from the intermembrane space into the cytoplasm [6,7]. Regulation of the progression towards apoptosis is directed by interactions on the OMM between three distinct subgroups of the BCL-2 family: the activator/sensitizer BH3 (BCL-2 homology 3)-only proteins, the pro-survival inhibitor proteins, and the pro-apoptotic executioner proteins [8,9]. Proteins of the BCL-2 family share amino acid sequences of homology known as BCL-2 homology (BH) motifs.

The pro-survival proteins (Bcl-2, Bcl-xL/Bcl2l1, Bcl-w/BCL2l2, Mcl-1, Bcl2a1/Bfl1, and Bcl2l10), along with the pro-apoptotic proteins (Bax, Bok, and Bak) share four BH motifs (BH1-4) [8,9]. They adopt a similar globular structure composed of nine α-helices, folding into a bundle, enclosing a central hydrophobic α-helix. This fold fosters a hydrophobic surface cleft, which constitutes an interface for the binding with BH3 motifs in other Bcl-2 family members. The globular multi-BH motif members of the Bcl-2 family mainly exert their apoptotic involvement at the OMM, to which they anchor by a C-terminal transmembrane region, exposing their globular helical bundle to the cytoplasm [10]. Unlike the globular Bcl-2 proteins, BH3-only proteins contain only one BH motif (BH3), which is often located in intrinsically disordered regions [11,12]. As many other IDPs, BH3-only protein folds upon binding, where their BH3 region becomes an amphipathic helix [12,13]. The BH3-only proteins can be divided into activators and sensitizers according to how they exploit their pro-apoptotic function [14,15]. Activators carry out their function by binding to pro-apoptotic proteins allowing the permeabilization of OMM and subsequent apoptosis event. Sensitizers bind to pro-survival members, inhibiting their binding with activator BH3 proteins and making them available to bind pro-apoptotic proteins.

Despite the importance of the BH3 motif in cell death regulation, a clear-cut definition of the motif is currently missing [16–18]. BH3-only proteins feature also distinct binding profiles and specificity toward Bcl-2 family members [19]. Attempts to define a consensus motif have returned motifs that are too strict (i.e., excluding proteins that are experimentally proved to bind Bcl-2 members) or too inclusive (i.e., reporting false-positives) [17]. A common feature, at the structural level, seems to be the presence of an amphipathic helix composing the BH3 motif that binds to the hydrophobic cleft on globular Bcl-2 members. This mainly happens by the insertion of four hydrophobic residues into hydrophobic pockets in the cleft and an invariant salt bridge between a conserved arginine residue in the globular Bcl-2 proteins and a conserved aspartate in the BH3-only protein [20]. One of the four hydrophobic residues, an invariant leucine, packs against and form interactions with conserved residues in the hydrophobic cleft of the globular Bcl-2 proteins [20].

One of the cancer hallmarks is the capability to escape programmed cell death, for example due to overexpression of pro-survival proteins of cancers [21–23]. Overexpression of pro-survival proteins is considered to contribute to tumorigenesis, the resistance of tumors to cytotoxic anticancer treatments, along with increased migratory and invasive potentials [24–26].

Due to their pivotal role as inhibitors of apoptosis, pro-survival Bcl-2 proteins have been amenable therapeutic targets for drug discovery. Advances in the knowledge of their interactions with BH3-only proteins at the structural level have led to the development of small molecules (BH3-mimetics), targeting the hydrophobic cleft and inhibiting the action of these proteins [27]. Some of these molecules although suffer of issues related to specificity and selectivity and are often working on specific cancer (sub)types [8]. To better exploit BH3-mimetics in cancer therapy, it is thus important to clearly elucidate the transcriptomic and mutational signature and abundance of the pro-survival proteins and their interactors in different cancer types, along with to identify the key interactions with their BH3-only modulators. For example, several studies have demonstrated how the affinity of pro-survival proteins to these mimetics varies to a large degree. Merino et al. [28] found that increased levels of Bcl-2 promote sensitivity to ABT-263, whereas increased levels of Bcl-xL or Bcl-w conferred resistance to the same mimetic. Similarly, two independent [29,30] found that cells overexpressing Bcl-2 showed high sensitivity to the BH3-mimetic ABT-737. In contrast, overexpression of Mcl-1 and Bcl2a1 in cell lines were found to confer resistance to a number of mimetics [30]. Comprehensive studies into the abundance, modifications and interactome of each of the pro-survival Bcl-2 members for each cancer type would generate important knowledge useful for optimization and design of BH3-mimetics. To achieve this goal a system-biology approach is needed and the integration of different layers of bioinformatic tools could help.

Apart from the deregulation of expression level of oncogenic and tumor suppressors observed in cancer, cancer develop when somatic mutations within the DNA alter specific amino acids in the protein-product, conferring selective advantages to highly proliferating cells [31]. Resistance to apoptosis is one of these selective advantages. Proteins are marginally stable under physiological conditions and the substitution of single amino acids can greatly alter their stability [32]. Besides protein stability, mutations can affect the binding affinity to interactors when occurring at, or close to binding sites [33,34] or even at distal site, through complex allosteric mechanisms [35,36]. The quantitative analysis of the effects of mutations on both the stability and binding affinity is thus crucial for understanding the capacity of the pro-survival proteins to propagate apoptosis. Another important aspect to consider in terms of effects induced by mutations of Bcl-2 proteins is that their mutations might alter the sensitivity to BH3-mimetics [14,37]. For example, Fresquet et al. [38] found that in BCL-2-expressing mouse lymphoma cells, two missense mutations within the BCL-2 gene confer resistance to the BH3-mimetic ABT-199. Despite the importance of pro-survival Bcl-2 proteins, no comprehensive studies have, to our knowledge, been designated to understand their molecular mechanisms in cancer by investigating the interface between transcriptomic, mutational signatures and the structural and functional effects of these alterations. Some of these aspects have been only analysed separately and on specific case study [22,39,40]. We here propose a “multiscale” approach to shed light on the pro-survival Bcl-2 proteins, by bridging two of the major branches of bioinformatics: (i) analysis of high-throughput sequencing data, and (ii) molecular modelling to unveil cancer-related alterations. We focused, as a case study, on Breast Invasive Carcinoma (BRCA) data from The Cancer Genome Atlas (TCGA) [41,42]. We identified a set of candidate genes encompassing Bcl-2 family members and their protein interaction partners containing the BH3 motif, revising its definition according to recent findings [17]. We then exploit differential expression analyses applied to RNA-Seq data from the TCGA-BRCA study to identify the expression levels of the candidate genes in tumor and normal tissue, along with in different breast cancer subtypes. We focused also on the alterations in terms of missense mutations altering the protein product in the same cancer (sub)type. Next, we zoomed in at the structural level integrating different computational methods, which allows assessing the impact on protein function or stability. As a result, we provide a comprehensive picture of the most important protein-protein interactions within the Bcl-2 family and their alterations in breast cancer or breast cancer subtypes, which can be used as a guide for future drug design or cancer target selection. Moreover, we predicted new BH3-only proteins interesting in a breast cancer context which would be amenable for future experimental research.

## Results

### Definition of the BH3 motif

Our consensus motif was defined, comparing previously described BH3 motifs [17,43,44], along with structural information on important residues for interactions in canonical BH3-only proteins [18]. We defined a ‘loose’ consensus motif with the goal to be permissive and avoid the loss of true-positives. A generalized motif composed of 10-13 residues was applied: <u>[ar,h,s]-</u>X(3,4)-<u>L</u>-X(2,3<u>)-[ar,h,s]</u>-[G,A,S,C]-X(0,1)-[D,E,Q,N]. *ar, h* and *s* stand for aromatic (W, Y or F), aliphatic hydrophobic (V, I, L, M), and small residues (A, C, P) respectively (**Figure 1**). As a consequence, the positions 1, 3, and 5 of the motifs are the hydrophobic residues for the *h1, h2* and *h3* hydrophobic pockets [8]. The leucine at position *h2* is very conserved [45], whereas the hydrophobic residues for the other binding pockets might vary in size and properties. The position 6, which was originally expected to require a glycine can also tolerate other residues with small side chains, such as alanine, serine or cysteine [17]. The position 8, which was originally expected to have an invariant aspartate, could tolerate mutations to glutamate, asparagine or glutamine, in light of sequences collected in a recent study [17] and reported BH3-like motifs of an autophagic protein [46,47].

**Figuer 1.**
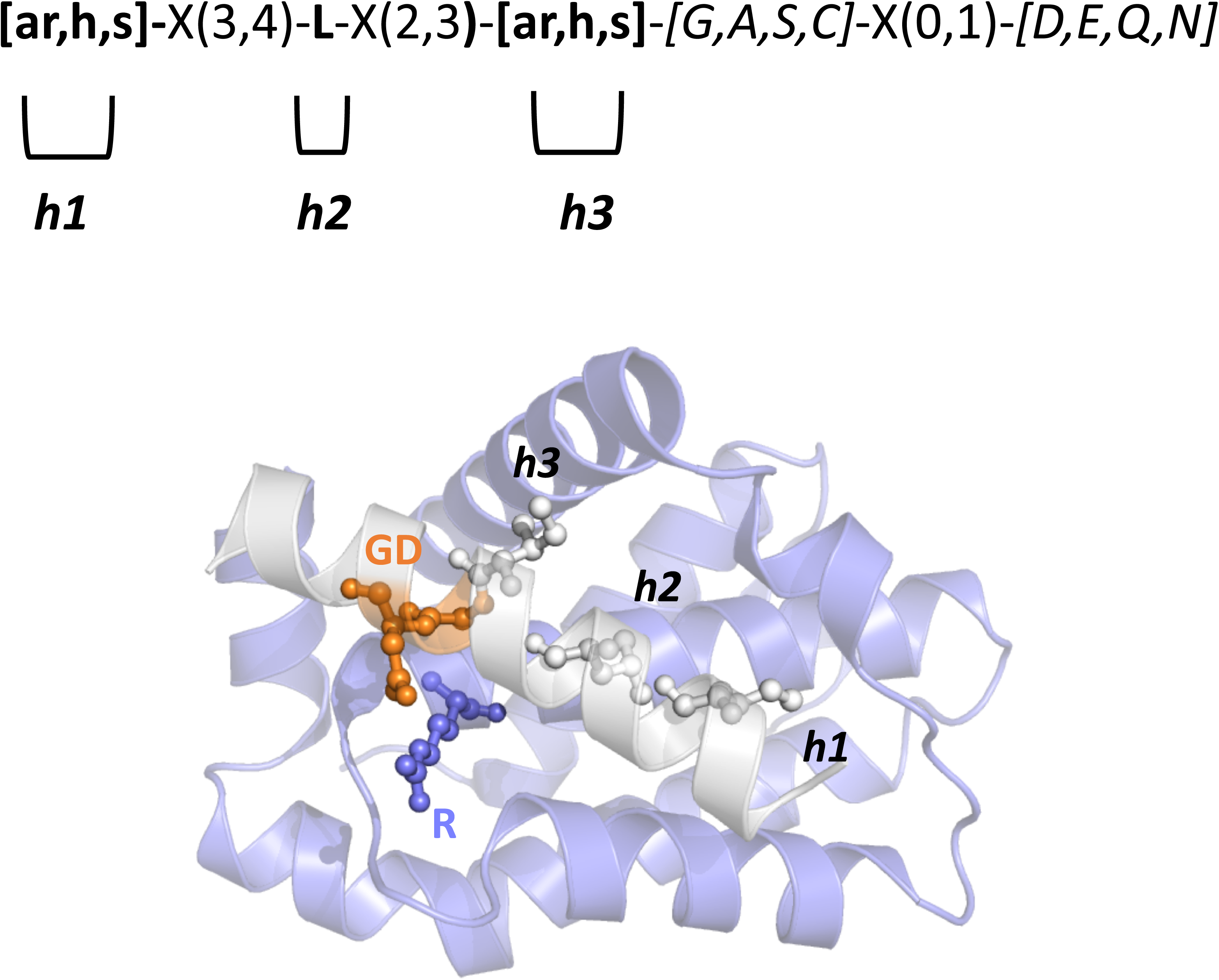
BH3 motif and key positions for Bcl-2/BH3 interaction. We illustrate the definition of the BH3 motif used in this study, highlighting the most important conserved residues for the binding to the BH3-binding groove of Bcl-2 family members. We used the complex of Bcl-xL and Bim as a reference (PDB entry 4QVF). The hydrophobic or aromatic residues which can occupy *h1* and *h2* hydrophobic pockets, along with the invariant leucine for h3 and the salt-bridge between the BH3 aspartate residue and the arginine of the Bcl-2 proteins are highlighted.

### Identification of BH3-containing interaction partners and BCL-2/BH3 interaction network

We retrieved the experimentally known BCL-2 family interaction partners from the human *Integrated Interaction Database (IID)* [48]. We then filtered the interaction list to retain only the proteins which included the BH3 motif. A total of 560 protein-protein interactions for the globular Bcl-2 members (Bcl-2, Bcl-xL/Bcl2l1, Bcl-w/Bcl2l2, Mcl-1, Bcl2l10, Bcl2A1, Bok, Bax, Bak, Bcl2l12, Bcl2l13, Bcl2l14, and Bcl2l15) were found (**Table S1**) and 295 of then were selected as possible BH3-containing proteins (**Table S2-S3, Figure 2**). Among the 295 proteins, 282 can be classified as BH3-only (**Table S2-S3**). The resulting protein-protein interaction network is compact and divided in only two connected components (**Figure 2**). The main connected component includes most of the Bcl-2 members and BH3-only proteins (291 nodes). An isolated small component refers to Bcl2l15 and its three BH3-only interactors (Meox2, Tead2, and Sdcbp). Most of the nodes feature a degree lower than 10 and collectively the average number of neighbors is 3.051. The most important hubs in the network (i.e., nodes connected to other nodes with a degree higher than the average connectivity in the network) are Bcl-2 (degree of 108), Bcl2l1 (96), Bax (88), Mcl-1 (62), Bak1 (33), Bcl2l2 (22), and Bcl2a1 (19) (**Figure 2A**). Most of these proteins also correspond to the ones with high values of closeness centrality, which measure important nodes for the network communication (**Figure 2B**).

**Figuer 2.**
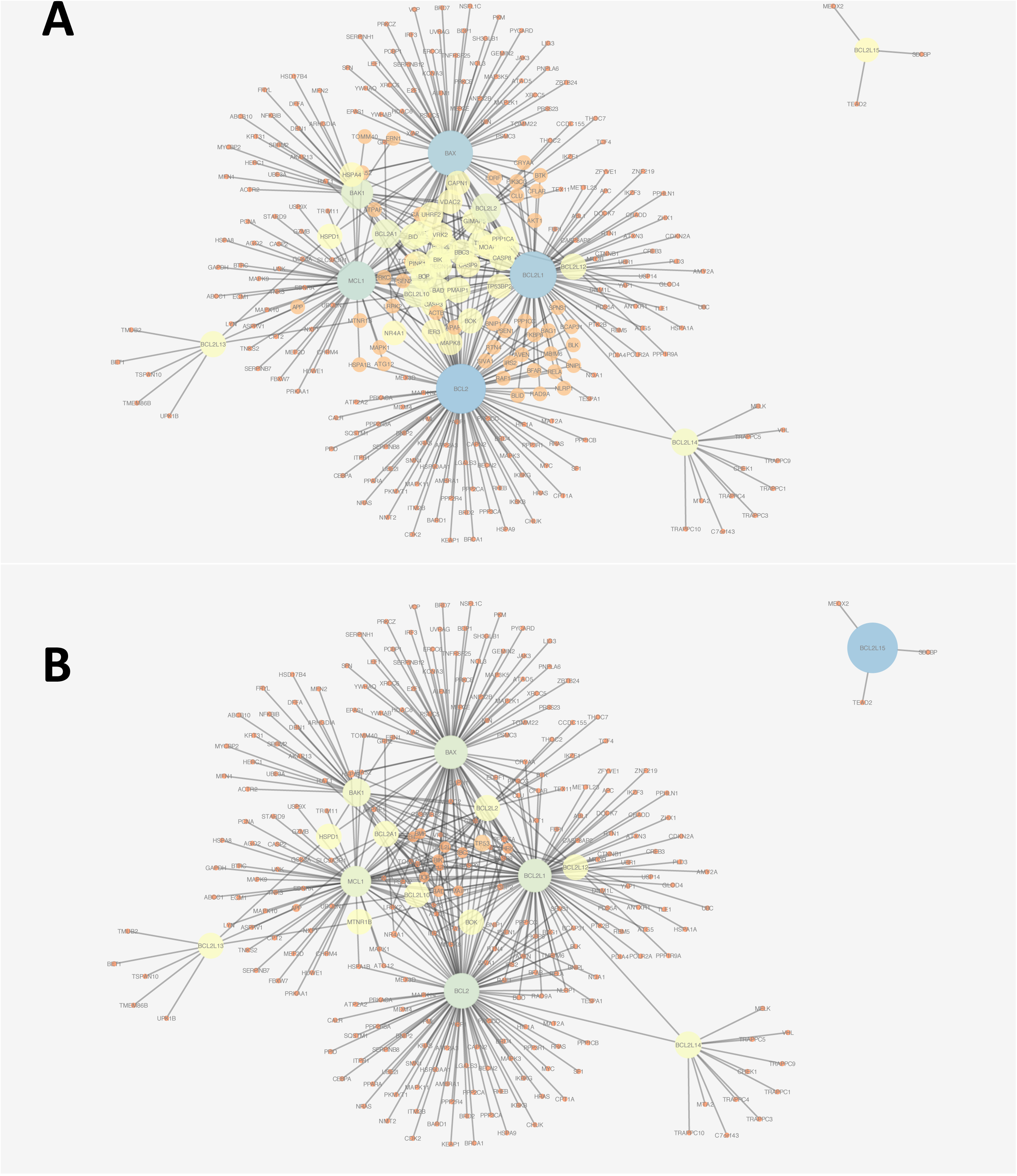
Protein-protein interaction network of Bcl-2 proteins and BH3-containing interactors in *IID*. The two connected components of the interaction network among the predicted BH3-only proteins and Bcl-2 pro-survival or pro-apoptotic globular proteins are shown. The nodes of the network are depicted with different size and shade of colors as a function of the degree (i.e., the number of edges for each node, A) and closeness centrality (B). It results that the most important hubs or nodes for network communications are the pro-survival Bcl-2, Bcl2l1, Mcl-1, Bcl2l2 and Bcl2a1, along with the pro-apoptotic Bax and Bak1.

We also compared our predictions to a manual curation from literature of experimentally validated BH3-only targets, where we identified 26 of our candidates as already known canonical or non-canonical BH3-containin proteins (Atg12, Aven, Beclin-1, Blid, Bnip1, Bnip2, Bop, Clu, Huwe1, Antxr1, Bbc3/Puma, Bcl2l11/Bim, Bad, Bid, Bik, Bmf, Hrk, Pmaip1/Noxa, Itm2b, Moap1, Rad9a, Spns1, Casp3, Pcna, Mycbp2 and Ambra1, **Table S2**). The remaining predicted BH3-only targets would require further verification upon analyses of 3D structures (or models), which the predicted motifs could fit the requirement for a BH3 motif (i.e., in disordered or solvent-exposed helical structures). Nevertheless, this list provides a rich source of information for experimental validation of new BH3-containing proteins.

### Changes in gene expression of Bcl-2 family members in breast cancer

At first, we aim to investigate which genes, encoding globular Bcl-2 members and their putative BH3-like interactors, are characterized by changes in expression levels, between the compared conditions: tumor tissue and tumor-adjacent normal tissue. The aggregated TCGA BRCA dataset originally contained 1102 tumor samples, and 113 tumor-adjacent normal tissues samples. Breast cancer is a heterogeneous disease at both the morphological and molecular level. To increase our understanding of biologically induced variation, we included PAM50 molecular subtype information, classifying breast carcinomas into subtypes based on variations in gene expression patterns. As a result, we excluded samples lacking subtype information. We performed differential expression analysis (DEA) on a final dataset, after pre-processing, normalization, and filtering, containing 14273 protein-coding genes and 444 tumor samples, including subtype information (Luminal A, Luminal B, Basal-like, and HER2-enriched) and 113 tumor-adjacent normal tissues samples. The Normal-like subtype was filtered out as it only encompassed four samples.

To quantify the magnitude and significance of differential expression between the conditions, tumor and normal samples, we employed *limma-voom* [49]. We identified 3092 differentially expressed genes, of which 1738 downregulated and 1354 upregulated in tumor compared to normal tissues. Among these genes, 45 candidate genes (BCL-2 family members or their BH3-only interactors) were differentially expressed with 21 of them upregulated and 24 downregulated (**Table S4, Table 1, Figure 3**). BCL2-A1 results the only pro-survival BCL-2 gene which is up-regulated in the majority of the comparisons. We also noticed that one of the main pro-survival genes, i.e. BCL-2 is mostly downregulated in the TCGA-BRCA samples with HER2 and basal subtypes.

**Table 1.** Differentially expressed BCL-2 genes in breast cancer subtypes. The logFC is indicated in the table, all the results refer to an FDR < 0.05. Empty lines indicated that the gene is not differentially expressed in the corresponding comparison. BCL2-A1 results the only pro-survival BCL-2 gene which is up-regulated in the majority of the comparisons. We also noticed that one of the main pro-survival genes, i.e. BCL-2 is mostly downregulated in the TCGA-BRCA samples with HER2 and basal subtypes. We did not report the comparison between LumA and LumB since it resulted in none of the BCL-2 family members with signs of deregulation.

| GENE | Cancer<br>vs<br>Normal | Basal<br>vs<br>HER2 | Basal<br>vs<br>LumA | Basal<br>vs<br>LumB | Basal<br>vs<br>Normal | HER2<br>vs<br>LumA | HER2<br>vs<br>LumB | HER2<br>vs<br>Normal | LumA<br>vs<br>Normal | LumB<br>vs<br>Normal |
| --- | --- | --- | --- | --- | --- | --- | --- | --- | --- | --- |
| BCL-2 |  |  | -2.35 | -1.91 | -2.20 | -2.26 | -1.82 | -2.11 |  |  |
| BCL2A1 | 1.31 | 1.25 | 2.02 | 1.61 | 2.77 |  |  | 1.51 |  | 1.15 |
| BCL2L1 |  |  |  |  |  |  |  |  |  |  |
| BCL2L2 | -1.01 |  |  |  | -1.17 |  |  | -1.21 |  |  |
| BCL2L10 |  |  |  |  |  |  |  |  |  |  |
| MCL-1 |  |  |  |  |  |  |  |  |  |  |
| BAK1 |  |  |  |  | 1.34 |  |  | 1.03 |  |  |
| BAX |  |  |  |  |  |  |  |  |  |  |
| BOX |  |  |  |  | -1.46 |  |  | -1.47 | -1.06 |  |

**Figuer 3.**
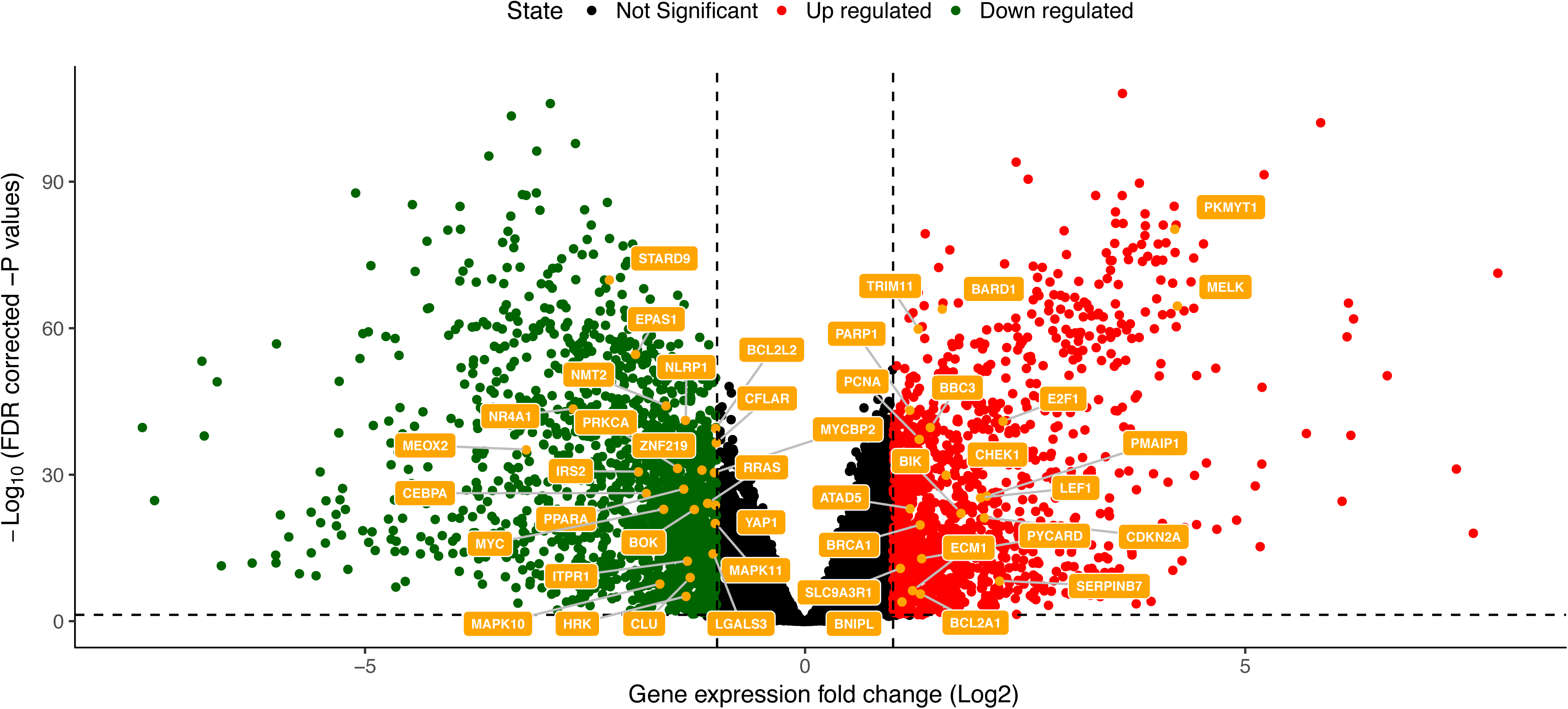
Differentially expressed BCL-2 and BH3-containing genes. We showed as a volcano plot the BCL-2 and BH3-only genes which are among the list of DE genes between tumor and normal tissues. The horizontal dashed line shows the FDR cutoff (0.05). The vertical dashed lines represent the cutoffs in terms of logFc for up- and down-regulated genes. The other comparisons at the subtype level are reported in the *Github* repository associated to the publication and summarized in Table S4.

We then analyzed the patterns of deregulation in the BH3-containing candidate genes (**Table S4, Table 2**). BH3-containing proteins are often induced transcriptionally by cytotoxic stresses and they can either inhibit pro-survival Bcl-2 proteins, or act through direct activation of pro-apoptotic Bak, Bax and Bok [8]. We thus were interested in pattern of opposite deregulation of BH3-only candidates with pro-survival Bcl-2 family members, respectively. In particular, in cancer samples evading cell death, we expect an up-regulation of pro-survival BCL-2 family members and down-regulation of the cognate BH3-only proteins. Another direction could be related to down-regulation of a pro-apoptotic BCL-2 member (BAX, BAK1 or BOK) and down-regulation of the cognate BH3-only. Only BOK is down-regulated in our dataset and we did not find any BOK-specific BH3-containing interactor deregulated. We thus focused on the relationship between the pro-survival and BH3-containing genes.

**Table 2.** Up-regulated BH3-only candidate genes in breast cancer subtypes. The logFC is indicated in the table, all the results refer to an FDR < 0.05. Empty lines indicated that the gene is not differentially expressed in the corresponding comparison. We did not report the comparison between LumA and LumB since it resulted in only three BH3-only genes with signs of deregulation (see Table S4). We reported only BH3-containing genes for which the differential expression was observed in at least two comparisons, for sake of clarity. The full list is reported in Table S4.

| GENE | Bcl-2 interactor | Cancer vs Normal | Basal vs HER2 | Basal vs LumA | Basal vs LumB | Basal vs Normal | HER2 vs LumA | HER2 vs LumB | HER2 vs Normal | LumA vs Normal | LumB vs Normal |
| --- | --- | --- | --- | --- | --- | --- | --- | --- | --- | --- | --- |
| RTN1 | Bcl2l1 |  |  | -1.65 | -1.13 | -2.12 | -1.44 |  | -1.91 |  |  |
| NMT2 | Bcl-2 | -1.57 | 1.14 | 1.06 | 1.38 |  |  |  | -1.82 | -1.74 | -2.06 |
| CLU | Bax, Bcl2l1 | -1.30 |  | -1.15 | -1.02 | -2.17 |  |  | -1.47 | -1.01 | -1.15 |
| ZNF219 | Bcl2l1 | -1.16 |  |  |  | -1.55 |  |  | -1.47 |  | -1.21 |
| ECM1 | Mcl-1 | 1.22 | -1.72 | -1.86 | -1.85 |  |  |  | 1.44 | 1.58 | 1.58 |
| SLC9A3R1 | Bcl2a1 | 1.08 | -1.42 | -1.48 | -2.17 |  |  |  | 1.22 | 1.28 | 1.97 |
| HRK | Bcl-2, Bcl2l1, Bcl2l2, Mcl1, Bcl2a1, Bax | -1.34 | 2.22 | 3.79 | 3.38 | -1.33 | 1.57 | 1.16 |  | -2.45 | -2.04 |
| IRS2 | Bcl-2, Bcl2l1 | -1.88 | 1.05 |  |  | -1.92 | -1.27 | -1.07 | -2.98 | -1.71 | -1.91 |
| NLRP1 | Bcl-2, Bcl2l1 | -1.35 |  |  |  | -1.27 |  |  | -1.43 | -1.28 | -1.65 |
| LGALS3 | Bcl-2 | -1.04 |  |  |  | -1.19 |  |  |  |  | -1.46 |
| RRAS | Bcl-2 | -1.10 |  |  |  | -1.17 |  |  |  | -1.01 | -1.39 |
| ITPR1 | Bcl-2 | -1.33 | -1.46 | -2.12 | -1.74 | -2.92 |  |  | -1.45 |  | -1.17 |
| CFLAR | Bcl2l1, Bax | -1.00 |  |  |  |  |  |  |  | -1.04 | -1.27 |
| NR4A1 | Bcl-2, Bcl2l10, Bcl2a1 | -2.63 |  |  |  | -2.64 |  |  | -2.72 | -2.55 | -2.83 |
| STARD9 | Mcl-1 | -2.21 |  |  |  | -2.10 |  |  | -2.50 | -2.16 | -2.43 |

Of note, the high overexpression of MELK (**Table S4**) in most of the comparison is likely due to effects unrelated to the apoptotic pathway, since this genes encodes for an oncogenic kinase in breast cancer [50]. Similarly, the up-regulation of PCNA, CHECK1 or GZMB (**Table S4**) could be more related to other aspects of apoptosis or breast cancer pathways. PCNA is known as a marker for breast cancer [51], even if this should be verified at the protein level, considering that we are analysing only gene expression data. Associations with CHECK1 levels and breast cancer have been also reported [52].

A group of predicted BH3-only, found in the interactome of at least one Bcl-2 pro-survival protein, are highly down-regulated in all or most of the breast cancer subtypes compared to the normal samples. It would be thus interesting to assess if the corresponding protein products could also bind and regulate other Bcl-2 family members which are prone to up-regulation in breast cancer. BH3-only genes with these patterns are, for example, CLU, IRS2, NLRP1, NMT2, ITPR1, CFLAR, LGALS3, RRAS, RTN1, STARD9 and ZNF219, which are interactors of Bcl-2 family members with minor deregulation in this study according to *IID*. Our results, also in light of the intrinsic incompleteness of the annotations in protein-protein interaction databases, suggest that they could be interesting candidates for future study to assess their promiscuity of binding towards other Bcl-2 proteins, such as Bcl2a.

SLC9A3R1 is down-regulated in the Basal with respect to the other subtypes (**Table 2**) and it is a Bcl2a1 interactor, which in turns is up-regulated in the same subtype (**Table 1**), suggesting an interesting association.

Of interest, we also observed that the pro-survival BCL2A1 and its interactor HRK (Harakiri) and were differentially expressed in the direction of an up-regulation of the pro-survival protein and a down-regulation of the inhibitor of apoptosis BH3-only HRK in all the subtypes with respect to the normal samples. Moreover, Hrk is a rather promiscuous BH3-containing interactor, considering that it has been found in the interactome of Bcl-2, Mcl-1, Bcl2l1, Bcl2l2 and Bcl2a1 (**Table 2**). Moreover, our analysis of the Bcl-2/BH3 protein-protein interaction network points out a high closeness centrality score (0.48) for Hrk (**Figure 2B**). Hrk has been experimentally showed to bind Bcl-2 and Bcl-xL [53]. Our results suggest that the interaction between Bcl2a1 and Hrk could be of interest to explore further in breast cancer considering that their deregulation is in the direction of evading apoptosis, a cancer hallmark.

Nr4a1 can also be an interesting BH3-candidate for Bcl2a1 in breast cancer for similar reasons (**Table 2**). A Nr4a1-derived peptide has been reported with the capability to convert Bcl-2 into a pro-apoptotic molecule [54,55]. Our predictions suggest that Nr4a1 could include two potential BH3 motifs (at positions 201-213 and 386-398), which if experimentally validated could open new direction toward a multifaceted regulatory role of Nr4a1/Nur77 on Bcl-2 proteins. The two BH3 motifs of Nr4a1 are both placed in disordered region and not buried to the solvent, according to the analysis of the 3D structure of the C-terminal domain of the protein (PDB entry 4RZF, residues 351-598) or through prediction of helical and β-sheet propensity with *FELLS* [56]. Thus, they could be in principle accessible for interaction with the Bcl-2 family members.

### Models of interaction between Bcl2a1/Bfl1 and the BH3-only interactors Hrk

Along with changes in expression levels, other alterations that could triggered evasion of apoptosis in cancer can be related to somatic mutations in the coding region of the Bcl-2 proteins and their interactors, which will exert an effect in the protein products of these genes. Before analyzing the TCGA-BRCA somatic missense mutations in the coding region of BCL2A1 and its interactors, we built a 3D structural model of their protein complexes.

Mutations can impact on a myriad of different aspects at the protein level and it thus becomes fundamental to be able to assess them at different levels, as we recently showed in other works [57,58]. The knowledge of the structure of the targets of interest is fundamental in this context. In the Protein Data Bank (PDB), the structures of Bcl2a1 in complex with the BH3 regions of Bim, Bak, Noxa, tBid and Puma were available.

We thus employed comparative modeling to derive models of the 3D structure of the Bcl2a1 complex with the BH3-like sequences of Hrk, similarly to what we did for other short linear motifs in complex with folded proteins [59]. We collected two different models for the interaction of the two BH3-only motifs that we found in Hrk (i.e., 28-50 and 63-85). We used the complex between Bcl2a1 and Puma as a template with a known X-ray structure (PDB entry 5UUL, [60]) since it is the Bcl2a1-BH3 complex with the best atomic resolution (1.33 Å) available s. As a result of the comparative modelling approach, our target BH3 peptides are assumed to interact in a similar manner to the template structure at the binding interface. This assumption is supported by the generally conserved helical conformation of BH3 motifs upon binding to the Bcl-2 family members [8]. The comparative modelling approach here used is convenient to scrutinize the effect of mutations at the binding interface in a high-throughput manner. However, one should consider that this approach could suffer of limitation in the description of fine structural details and interactions that other more computationally-demanding and accurate methodologies could provide, such as sampling based on Montecarlo or Molecular Dynamics simulations.

**Figure 4A-B** shows the BH3 peptides of Hrk_1 compared to Puma (residues 132-154). The BH3 motif for Hrk_1 (28-50), as found in our motif search, defines the first residue as an aliphatic hydrophobic residue (L32 of Hrk_1). Nevertheless, Barrera-Vilarmau et al. [61] resolved a fragment of Hrk (residues 22-53, PDB entry 2L58) by NMR. They identified T33 opposed to L32 as the key residue in binding of Hrk with Bcl-2 and Bcl-xL. We thus modeled the complex Bcl2a1-Hrk_1 (28-50) aligning T33 as the binding residue in *h1.* The complex features hydrophobic residues at the positions 2,3 and 5 of the motif, with the invariant leucine in position 2 and the conserved aspartate occupying position 4 (**Figure 4B)**.

**Figuer 4.**
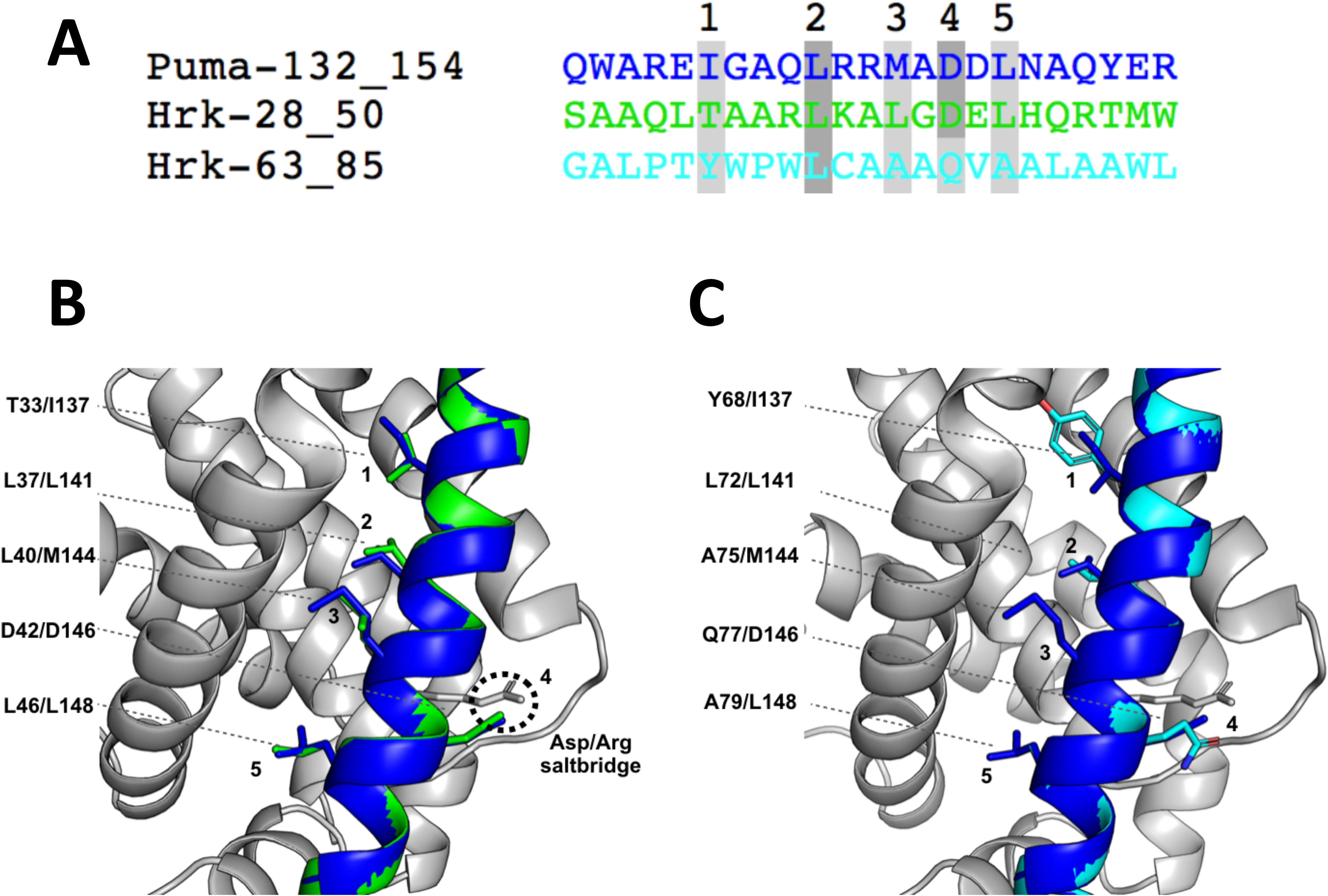
The model of interaction of Bcl2a1 and Hrk BH3-like peptides. A) The predicted BH3 regions of Hrk (i.e., Hrk_1, residues 28-50 and Hrk_2, residues 63-85) are aligned to the Puma BH3 as a reference. B-C) The 3D models of the complexes between Bcl2a1 and Hrk_1 or Hrk_2 are shown with the residues important for the intermolecular interactions, in comparison with the experimentally-derived complex.

A number of BH3-only proteins have been predicted to contain a transmembrane (TM) region, suggesting an outer mitochondrial membrane (OMM) anchoring function and in some cases the BH3 itself can associate to membranes [62]. Barrera-Vilarmau et al. revealed a transmembrane region in Hrk (residues 69-91) [61]. Given these results, we expect this second BH3-containing peptide (Hrk_2) to be part of the TM region and have a role in membrane targeting activities. Nevertheless, its association with membranes could be modulated by many factors and it could also act as interactor for the Bcl-2 family members when it is not associated to the membrane. Also, in this case we could identify the hydrophobic residues at position 1, 2, 3, and 5. Position 4 is characterized by a glutamine (Q77) residue, which replaces the negatively charged aspartate (**Figure 4A and C**). Despite aspartate being a highly conserved key residue in the BH3 motif, other residues at this position have been reported in BH3-like motifs [17,47]. For example, the pro-autophagic protein Ambra1, which acts as an inhibitor of pro-survival Bcl-2 through a BH3-like motif contains a glutamine residue instead of the conserved aspartate [47]. Another example is found in the BNIP group of proteins, a relative new subclass of BH3 proteins, where the aspartate is substituted by an asparagine [63]. Our structural analysis of Bcl2a1 /Hrk_2 (residues 69-91) complex, along with the other BH3-like sequences mentioned above, suggests that the invariant salt-bridge between the BH3-only negatively charged aspartate and the positively charged arginine of the Bcl-2 proteins, could be also replaced by interactions between the arginine and residues such as asparagine or glutamine, which are still capable of providing a delocalized partial charge around their functional groups.

### Structure-based assessment of the effect of cancer-related mutations in the Bcl2a1-BH3 complexes: stability and local effects on interactions

We collected missense mutations for the proteins of interest in breast cancer, aggregating data from different cancer genomics projects (see Materials and Methods for details and **Table S5**). Four mutations were reported for Bcl2a1 (M75R, L99R, Y120C and V145L), whereas we did not identify missense mutations altering Hrk in the breast cancer samples under investigation. We used the models of the two complexes of Bcl2a1 and Hrk BH3 regions, along with the structure of the complex between Bcl2a1 and Puma to predict the functional impact of any possible substitutions of the Bcl2a1 protein and its BH3-containing interactor on the binding free energies (**Figure 5**). We also used the same high-throughput pipeline to estimate the changes in free energies upon mutation associated to the structural stability of Bcl2a1, to be able to discriminate between effects that are related to its cellular function (i.e. binding with the BH3-only proteins) or related to its stability, and thus likely to alter for example the protein turnover at the cellular level (see below).

**Figuer 5.**
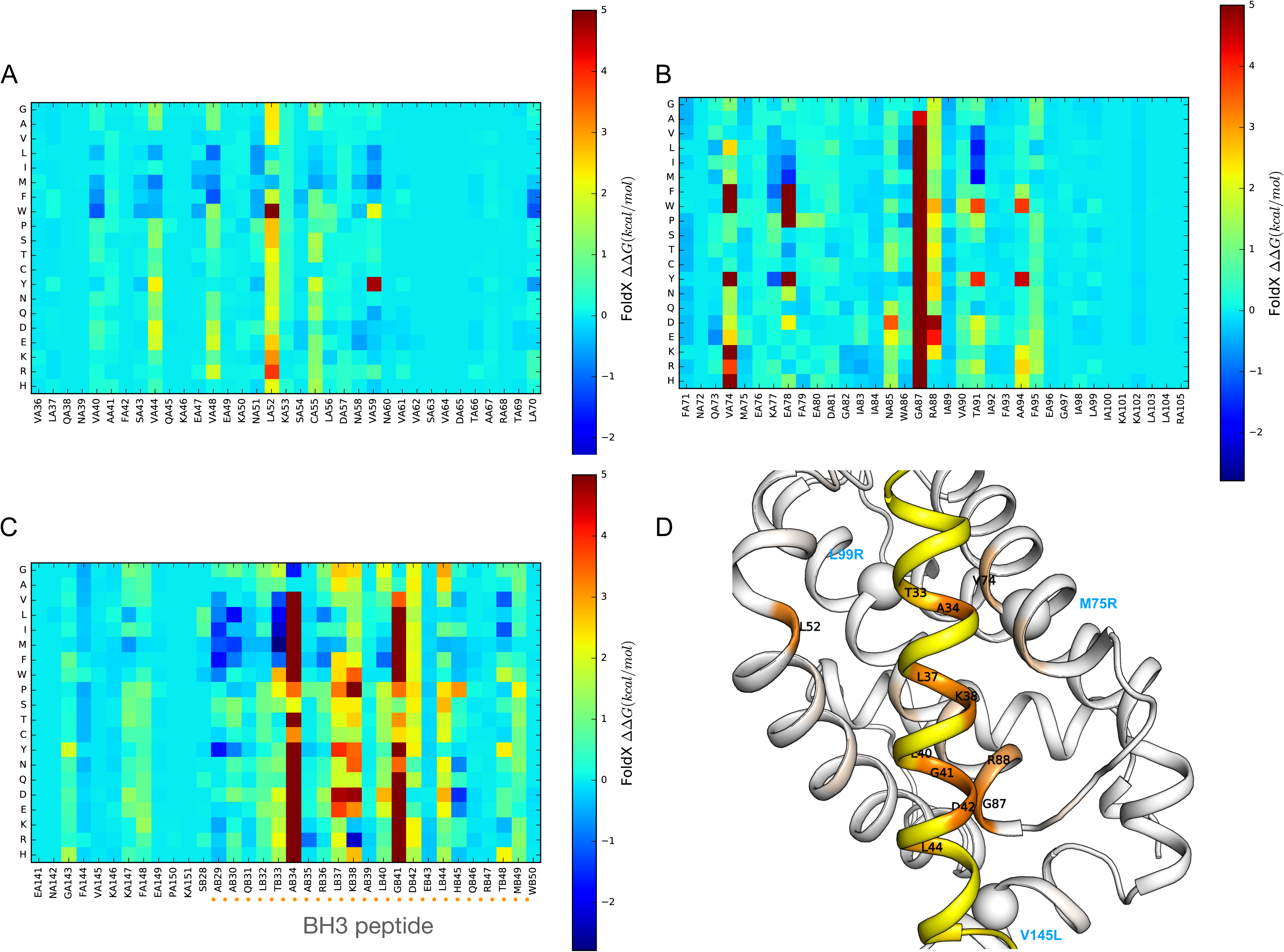
*In silico* deep mutational scanning of the Bcl2a1-Hrk complexes using an empirical energy function. The binding ΔΔ Gs are shown as heatmaps for the saturation mutagenesis carried out in silico on the Bcl2a1-Hrk_1 complex as a reference. In particular, the results for: A) α-helices 2 (residues 32-52), and 3 (residues 53-58) of Bcl2a1, B) α-helices 4 (66-80) and 5 (86-105) of Bcl2a1 and, C) Hrk_1 BH3 peptide (residue 28-50) are shown. The Δ ΔGs have been truncated at a value of 5 kcal/mol for sake of clarity. D) The color gradient indicates the average Δ Δ Gs for mutations at each position in Bcl2a1 (white to red) and Hrk_1 (yellow to orange) on the 3D structure. The spheres indicate positions of three of the breast cancer mutations of Bcl2a1 (M75R, L99R and V145L). The data with the mutational scans of each complex are reported in the Github repository.

In our mutational scans, ΔΔG values that do not change upon mutation or are close to 0 kcal/mol indicates that the original residue is not essential for protein stability and/or complex formation. Negative ΔΔG values indicate that the mutant is more stable than the wild-type variant, whereas positive ΔΔG values indicates that the mutation has a destabilising effect and that the wild-type amino acid may have an important function preserving the structural integrity of the protein or of the complex.

The high-throughput mutational approach allowed us to evaluate the effects, on Bcl2a1stability or interaction with BH3-containing proteins, of substituting wild-type residues with the mutant variants found in cancer samples, thus allowing a classification of potential damaging and neutral cancer mutations. Moreover, it allows us to evaluate the general effects of any amino-acid substitutions over the whole protein structure or complex, providing a useful set of precomputed ΔΔGs for future assessment or annotation of Bcl2a1 mutations. The latter could be a valuable source of information for future studies.

Additionally, the deep mutational scan increased our knowledge of hotspot residues for protein-peptide binding between Bcl2a1 and putative BH3-like proteins, which could potentially aid in the design of selective peptide inhibitors targeting Bcl2a1.

We identified R88 of Bcl2a1, which is the arginine important for salt-bridge interactions with the conserved aspartate of the BH3 motif, as a sensitive hotspot for mutations (**Figure 5B**). This position cannot tolerate any substitution, even to lysine. The only tolerated substitution is to histidine. The histidine side-chain has a pKa close to physiological pH, implicating that the local amino acidic environment determined shifts in pH, will change its average charge. Consequently, we could expect a population of protonated positively charged histidines in the complexes at physiological conditions. Other critical positions in the BH3-binding cleft of Bcla1 are hydrophobic residues such as L52 (**Figure 5A**) and V74 (**Figure 5B**).

T33 of Hrk, the position earlier identified as one of the residues contributing to the binding in the BH3 motif of Hrk, is predicted to favor substitution to a hydrophobic amino acid in the form of either: one of the aliphatic amino acids (alanine, valine, leucine, and methionine) or the aromatic phenylalanine (**Figure 5C**). This result suggests that T in *h1* is suboptimal for binding to Bcl2a1 and this feature could be used to design a stronger binder. A34 and G41 of the BH3 peptide are predicted highly intolerant to any substitution except for a glycine and alanine, respectively. This suggest that the two BH3 positions adjacent to the residues for *h1* and *h2* binding need to be of a small size and any other substitution would create a steric hindrance. We observed a similar trend in the mutational scan of the complex between Bcl2a1 and Puma (**Figure S1**), suggesting important features to define the BH3 motif.

The BH3 motif invariant leucine L37 in *h2* is predicted sensitive to most substitutions. L37 moderately tolerates substitutions by valine, isoleucine, and methionine. L40 (*h3*) and L44 (occupying a fourth hydrophobic pocket, *h4*) also contribute to the complex formation and are predicted to tolerate substitutions only to aliphatic and aromatic amino acids.

The mutation of the conserved and salt-bridge forming aspartate (D42) is generally poorly tolerated. D42 can be substituted without any marked effects only to: (i) glutamate, which is also negatively charged (ii) asparagine and glutamine, which are similar in size to aspartate and glutamate but contain an amino group. This finding consolidates the notion that the selectivity at this BH3 site is triggered by the possibility of maintaining electrostatic-based interactions with the arginine of the BCL-2 protein. This can occur also in absence of negatively charged residues, if polar residues of a similar size are present, such as glutamine and asparagine. The original definition of the BH3 motif, which expected an invariant aspartate, should thus be revised to include this possibility so that a larger number of interactors can be identified.

Conclusively, the mutational scan of Bcl2a1-Hrk_1 (residues 28-50) pointed to a subset of substitutions with damaging effects on the complex formations. These results suggest the importance of: i) hydrophobic residues in binding the amphipathic helix comprising the BH3 motif; ii) propensity to small side chains adjacent to the *h2* and *h3* hydrophobic residues; iii) the disadvantage in terms of binding affinity of having a threonine (T33) compared to a hydrophobic amino acid in the *h1* pocket; and iv) the electrostatic interactions between arginine (R88) and the conserved aspartate (DB42), which can be fulfilled by other residues such as glu, gln and asn.

### Structure-based assessment of the effect of cancer-related mutations in the Bcl2a1-BH3 complexes using protein structure networks

The four reported cancer mutations in Bcl2a1, L99R, Y120C and V145L are not located in the proximity of the BH3-binding domain and are not predicted to have any local effects on the complex formation (**Figure 5 and Figure S1**). M75 is in proximity of the BH3 binding pocket and it has been mentioned as important residue in the hydrophobic pocket *h2* [64]. M75R is not predicted to alter binding ΔΔGs in our analyses, in agreement with the fact that most of the interaction with the BH3 peptide could be mediated by the backbone of this residue [65].

We thus investigate possible allosteric effects induced from these distal sites to the BH3 interface using a contact-based Protein Structure Network (PSN) approach [66,67]. Indeed, indirect and long-range effects of the mutations to the BH3-binding site cannot be captured by the high-throughput mutagenesis scan that we performed with the *FoldX* empirical energy function, which is tailored to describe local rearrangements at the side-chain level only.

At first, we generated an ensemble of conformations (**Figure 6A**) around the structure of a reference Bcl2a1 -BH3 complex (see Materials and Methods for details) using a coarse-grained model and the *CABS-flex* sampling method [68,69]. The models were then reconstructed to all-atom representation before the PSN analyses. We used an ensemble of conformation for better model the inherent flexibility of the proteins in the complex, along with to remove spurious contacts from the PSN. We then analyzed the Bcl2a1 cancer mutation sites for their: i) propensity to act as a hub-residue in the network, i.e. a node that features a high degree of connectivity and thus likely to be important for the maintenance of the protein architecture or for communication throughout the network; ii) propensity to communicate to the BH3-binding region, estimating the shortest paths of communication between each mutation site and a group of residues (V48, R88, L52, V74, T91, and F95, **Figure 6B**), which we selected as hotspots for binding by the deep mutational scan discussed above (**Figure S1**).

**Figuer 6.**
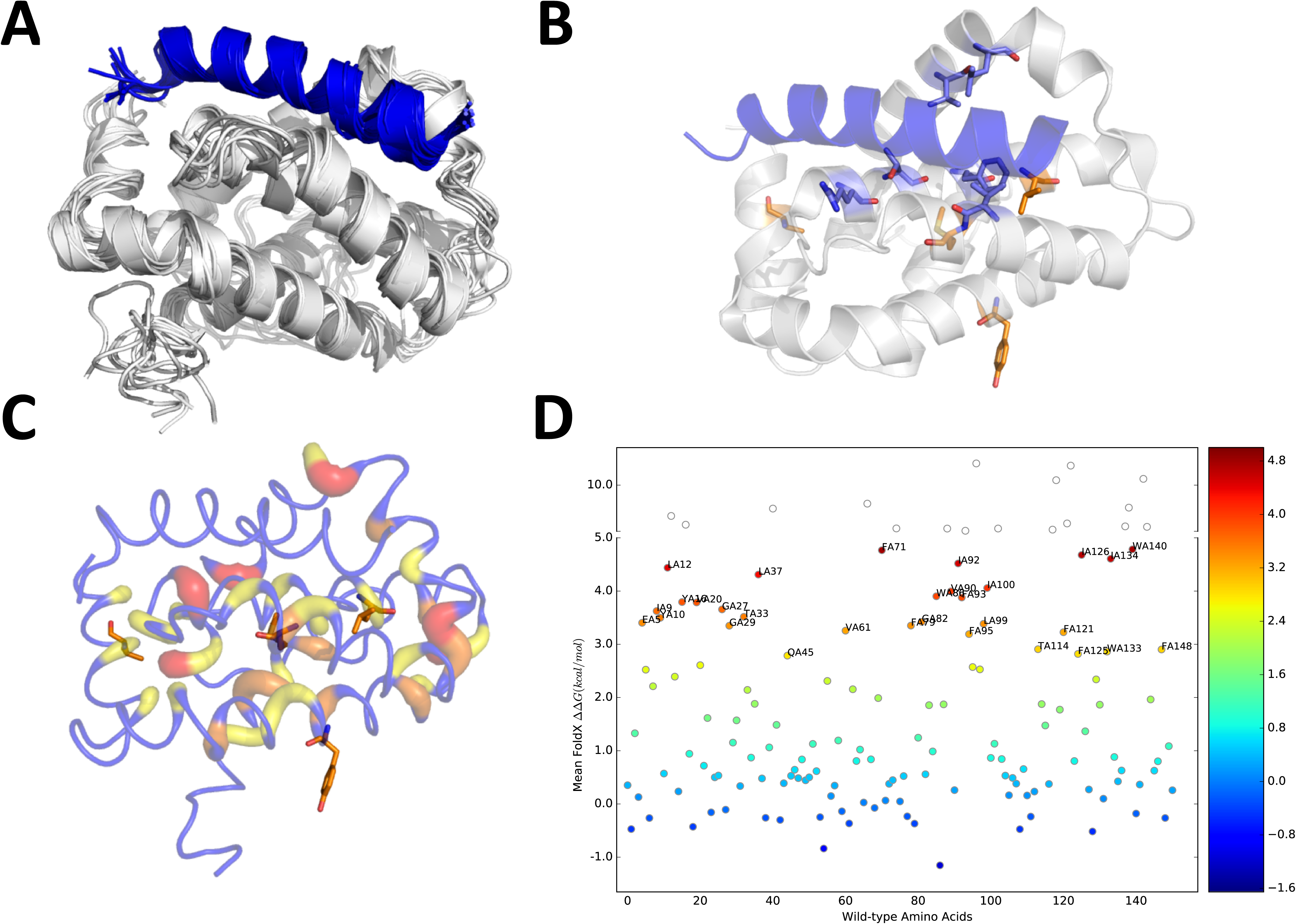
Analysis of the mutation sites in light of PSN of a conformational ensemble of Bcl2a1-Puma complex. **A)** The ten models of the conformational ensemble of the Bcl2a1-Puma complex generated by *CabsFlex 2.0* is shown. We used this ensemble of conformations for the PSN analysis. **B)** The Bcl2a1 cancer mutation sites and the target residues for path analyses (V48, R88, L52, V74, T91, and F95) are highlighted in orange and marine, respectively using as a reference the X-ray structure of the complex between Bcl2a1 and Puma (PDB entry 5UUL) **C)** The hub residues in the PSN of the Bcl2a1-Puma complex are showed with different scale of colors and cartoon thickness as a function of the degree (from yellow tored for degree from 3 to 5). The nodes that are not classified as hubs (degree < 3) are colored in blue. M75 and L99 are hub residues in the PSN, whereas Y120 and V145 do not show a hub behavior. **D)** Effects of amino acid substitutions on the free-state of Bcl2a1 upon *in silico* saturation mutagenesis to estimate Δ ΔGs associated to protein structural stability. A scattered plot depicting the mean Δ ΔG of all the possible mutations in each position of the wild-type sequence of Bc2a1 is shown. The top 20 most destabilizing mutations are labelled. The labels follow the convention: residue type, chain ID, and residue number. L99 is among the top 20 hotspots for protein stability, suggesting its sensitivity to any kind of mutations.

M75 and L99 are hub residues in the PSN, whereas Y120 and V145 do not show a hub behavior (**Figure 6C**). Y120 is solvent exposed in the ensemble and thus unlikely to contribute with intramolecular interactions. V145, on the contrary, is partially buried in the protein core and next to a hub residue (F144).

As a complement to the hub analyses, we carried out the saturation scan on the free-state of BCL2-A1 with the empirical energy function described above (**Figure 6D, Table S6)**. This allowed us to predict the impacts of the cancer mutations on the protein structural stability, which were in overall agreement with the hub results. Indeed, we identified M75R and L99R as damaging mutations for stability (ΔΔ G of 3.32 and 6.33 kcal/mol, respectively). Y120C and V145L had neutral effects. L99 is also one of the BCL2-A1 hotspots upon the deep mutational scan (**Figure 6D**), suggesting the sensitivity of this site to mutations and thus its importance for the Bc2a1 architecture. In support to these results, M75 and L99 are also highly conserved sites in 137 Bcl2a1 homolog protein sequences according to a *ConSurf* analysis (conservation scores of −0.952 and −0.761, respectively, **Table S7**), whereas Y120 and V145 are poorly conserved.

Our analyses suggest that M75R and L99R mutations have the major effect of destabilizing the protein structure and could be for example validated assessing the protein cellular levels of this variant and its propensity for increased degradation by the proteasome, as we recently did for other cancer-mutations [70,71].

To assess the capability of the mutation sites to mediate long-range effects to the BH3 binding region we then calculated the shortest paths of communication between each mutation site and the interface residue probes depicted in **Figure 6B**. We manually discarded paths that were not likely to act through a cascade of collisional events mediated by changes in rotameric states of the interest residues, to reduce the risk of false positive hits. Y120 turned out to be a possible residue that from an accessible surface can communicate to three of the interface residues selected as probes in the path analyses, i.e. R88, V74 and F95 (**Table 3**). This communication pass through a conserved group of nodes around Y120 (E124, M127 and I9) in all the cases.

**Table 3.** Shortest paths of communication from Y120 to the BH3-binding interface of Bcl2a1.

| Target interface residue | Path | Path length | Sum of weights | Average weight |
| --- | --- | --- | --- | --- |
| **VAL74** | Y120-124E-127M-9I-123A-13A-122V-71F-75M-74V | 10 | 480 | 53.3 |
| **ARG88** | Y120-124E-127M-9I-126I-130T-84I-129N-79F-78E-88R | 11 | 510 | 51 |
| **PHE95** | Y120-124E-127M-9I-123A-13A-122V-71F-75M-74V-F95 | 11 | 580 | 58 |

## Discussion

The network of protein-protein interactions between globular Bcl-2 family members and their BH3-only interactors plays an important role in controlling tissue homeostasis and deregulation can lead to cancer development. An increased understanding, through different tools, of the alterations of Bcl-2 members and their network of interactions in cancer could be useful to better exploit them as therapeutic targets. Through this study, we presented a computational workflow aimed at providing insight into: (i) the expression of pro-survival Bcl-2 members and their interactors in a certain cancer type, (ii) elucidating the functional interaction between them, and predicting the effects of substitutions on these interactions, and (iii) identifying alterations which could impact on the turnover of the protein, altering its structural stability. We applied this workflow on the BRCA data from The Cancer Genome Atlas (TCGA), as an example. Our framework can be extended to any other cancer datasets deposited in GDC or data from similar genomic initiatives.

Despite the importance of the Bcl-2 family conserved BH3 motif in mediating protein-protein interactions, a BH3 consensus motif is elusive [16]. We defined the motif for our search of candidate interactors in light of literature reports and a recent work in which the BH3 motif has been redefined as a short linear motif [17,43,44]. We allowed a certain degree of flexibility in certain conserved positions to have a broader coverage and prevent the removal, in our search, of possible non-conventional BH3-like proteins. The motif was applied to filter interaction partner of the Bcl-2 family members, extracted from an integrated curation of protein-protein interactions [47], providing a collection of more than 250 possible BH3-containing proteins of which 26 have been already experimentally validated. The remaining candidates could be interesting targets for experimental validation upon verification that they are in disordered regions or exposed helical regions of the corresponding proteins, a requirement for a BH3-like regions.

The pro-survival members were shown to be upregulated in a variety of tumor types [21,22] and have additionally been considered to contribute to tumorigenesis and therapeutic resistance [25,26]. Further studies, with bioinformatic approaches, into the alterations of the pro-survival members at the mRNA and protein level in specific cancer (sub)types can thus generate knowledge to guide and optimize the anti-cancer treatments. This point is especially critical if we consider that not necessarily the same Bcl-2 proteins and BH3-containing interactors will be the fundamental ones to target for all the cancer types and that they have been shown to be able to compensate each other and contribute in this way to resistance to BH3 mimetics [8]. Here, from the analysis of the TCGA-BRCA dataset, we uncovered the gene expression landscape of globular Bcl-2 members and their putative BH3-like interactors in breast cancer. We found a major signature for the pro-survival BCL2A1 gene, which is upregulated in breast cancer and its subtypes. The expression and function of pro-survival BC2LA1 in normal tissues appears to be linked to the immune system in which, it seems that development of inflammasomes increases the expression of BCL2A1, consequently protecting pro-inflammatory cells from apoptosis [72]. Moreover, a physiological function of Bcl2a1 in the mammary glands have been identified, in which overexpression of BCL2A1 in mice has been linked to the prevention of mammary gland involution by apoptosis [73]. Moreover, a study comparing a range of solid tumor tissues found the highest expression of BCL2A1 in breast cancers [22], while another study comparing expression levels between stages of breast cancer found an association with worst survival of the patient and high expression levels of BCL2A1 [74]. Apart from the likely role in tumorigenesis, Bcl2a1 induces chemotherapeutic resistance by suppressing apoptosis upon toxic stimulation, consequently preventing cell death. Overexpression of BCL2A1 in cell lines has been found to promote resistance to different cancer drugs including the BH3 mimetic ABT-737, a specific inhibitor targeting Bcl-2, Bcl-xL, and Bcl-w [30]. Due to their significant role as inhibitors of apoptosis, pro-survival Bcl-2 proteins have been considered promising targets for anti-cancer therapy. Progress has been made and several BH3 mimetics targeting the hydrophobic cleft in pro-survival members, show promising perspectives [75–77]. What is common for these mimetics is that they have been successful in inhibiting Bcl-2, Bcl-xL, and Bcl-w, but not Bcl2a1, or they have been broad-spectrum inhibitors with differing affinities depending on the pro-survival target proteins. In spite of the general progress, no potent and selective BH3 mimetics, targeting Bcl2A1 has so far been demonstrated [27]. Unraveling to what extent putative BH3-like interactors are deregulated in breast cancer and additionally clarifying their possible interaction with Bcl2a1 at the structural level, might provide a valuable source of information. Identifying possible Bcl2a1 selective interactors could serve as templates for the design of BH3 mimetics, targeting and preventing its pro-survival role in tumors. An interesting approach to drug design for this protein has been recently proposed and can benefit from further knowledge on the interactome and specificity towards this underappreciated Bcl-2 family member [64].

We here identified two putative BH3-containing interactors of interest in the context of Bcl2a1, i.e. Hrk and Nr4a1, which are downregulated in the TCGA-BRCA dataset and in different BRCA subtypes, accompanied by upregulation of Bcl2a1, suggesting a signature of cell death evasion, which is not compensated by changes in the pro-apoptotic Bcl-2 family members. Another interesting predicted BH3-only interactor along this direction is Slc9A3r1 in the BRCA Basal subtype. Hrk has been already reported as a BH3-containing protein and its interaction with other Bcl-2 family members has been addressed experimentally [53,61]. Our study suggests to explore more extensively its interaction and cellular role with attention also to Bcl2a1. On the other side, Nr4a1 would need to be studied as a possible new and non-canonical BH3-containing protein.

We also provide a model of interaction for Bcl2a1 and the two BH3-like motifs of Hrk and a deep mutational scanning to identify the possible molecular determinants of their binding mode. One of the two BH3 motifs that we predicted for Hrk is presumable located in a transmembrane region [61]. We speculate that this could act as a ‘conditional’ BH3, which might be a sensor for structural changes induced by cellular conditions that can dissociate Hrk from the membrane and then bind to Bcl-2 family members in the BH3-binding groove. Experiments addressing this hypothesis could shed new light on the Hrk mechanism of action and rule out that the motif predicted by our study is not a false positive.

A proper assessment of the impacts of mutations on both the stability of Bcl-2 family members and of the binding affinity to BH3 interactors, is critical to understand the functional capacity of pro-survival proteins to propagate apoptosis. Drug resistance in anti-cancer treatments continues to be one of the leading reasons for unsuccessful treatments and several studies have linked mutations in Bcl-2 family members to altered sensitivity or resistance to BH3-mimetics [14,37,38]. In general, a structure-based functional and stability assessment of mutational data have lagged behind the growth of data generated from modern high-throughput techniques. Here, we applied a high-throughput deep mutational scanning to predict the effects of mutations found in breast cancer samples, on both the structural stability of Bcl2a1, and the binding free energies between Bcl2a1 and the BH3-only interactors. This high-throughput approach permitted us also to evaluate the general effects of any amino-acid substitution on stability and binding and suggest important positions of the BH3 or of the Bcl2a1 protein for the interaction. For example, we shed light on the requirement for small side chain residues in proximity of the hydrophobic residues for interaction with h1 and h3 hydrophobic pockets of Bcl2a1, along with the possibility to replace the conserved negatively charged residue of the BH3 motif with also the cognate polar residues, asparagine and glutamine. Moreover, we showed how the Thr occupying one of the hydrophobic pockets is suboptimal for binding and this knowledge could be exploited to design peptides with higher affinities at this position.

The deep mutational scanning allowed us to provide a more comprehensive view on the alterations of Bcl2a1 in breast cancer, beyond mere changes in expression levels. The Bcl2a1 mutations in breast cancer were not predicted to change locally the binding affinity but they showed, in two cases (L99R and M75R), a marked impact on protein stability suggesting that despite upregulation these specific variants, in some of the samples, could be compromised due to increased turnover. This result clearly shows how important is to account, in analyzing cancer alterations of a certain group of genes, also for the compensatory effects that can be produced by different layers of modifications occurring at the same time in a sample. At last, using a protein structure network approach, we have been able to identify a mutation site (Y120) which might trigger allosteric effects to the BH3 binding groove and, as a such, be a long-range modulator of the Bcl2a1 protein.

## Conclusions

In summary, we here proposed a ‘multiscale’ approach to unveil the pro-survival Bcl-2 signature in cancer, bridging changes in the expression levels across the whole protein-protein network and mutations of the pro-survival Bcl-2 members and their putative BH3-like interactors with changes at the structural level. We provided a computational workflow to uncover the gene expression landscape of the complex protein-protein interaction network for regulation of Bcl-2 family members, along with to analyze the structures of these complexes and the impact of mutations. Moreover, we have proposed a high-throughput *in silico* mutagenesis approach to identify functionally important residues in the pro-survival members and their interactors. Our study highlights the prospects of an integrative bioinformatic approach, as it could potentially aid the development of targets for development of new BH3 mimetics and at the same time identifying substitutions in pro-survival BH3-only interactors that would reduce binding to other pro-survival members without substantially weakening the binding to the selected target. Lastly, we note that the approach to pro-survival proteins, proposed here, can be applied to anti-apoptotic members as well. For example, the evaluation of cancer mutation in terms of classifying damaging or neutral mutations, would be also relevant with a focus on anti-apoptotic members.

## Materials and Methods

To reproduce this study, we released a Github repository where data, scripts, and guidelines are deposited (https://github.com/ELELAB/bcl_bh3only_breast_cancer).

### Identification of BH3 motif containing interaction partners

We used, a source of protein-protein interactions, the *Integrated Interactions Database* (*IID*) [78] of tissue and organism-specific interactions, downloaded on February 6th, 2018 (version 2017-04), to retrieve known interactions partners of the globular Bcl-2 family members (Uniprot identifiers: Bcl-2; P10415, Bcl-xL; Q07817, Bcl-w; Q92843, Mcl-1; Q07820, Bcl2-l10; Q9HD36, Bcl2a1; Q16548, Bok; Q9UMX3, Bax; Q07812, Bak; Q16611, Bcl2l12; Q9HB09, Bcl2l13; Q9BXK5, Bcl2l14; Q9BZR8, and Bcl2l15; Q5TBC7) in human tissues. Subsequently, we filtered the interaction partners to retain only those containing the definition of a consensus BH3 motif described in the results. The protein-protein interactions were visualized and analyzed as a network using *Cytoscape* [79]. Upon consultation of the recently released *BCL2DB* database [80], we discovered that some of the Bcl-2 family members are better classified as Bid-like proteins (i.e., Bcl2l12-15) and we discarded them from the following analyses.

### Analysis of TCGA-BRCA RNA-seq data

For this study, we aggregated RNA-Seq BRCA data from The Cancer Genome Atlas (TCGA), using the *TCGAbiolinks* Bioconductor package v 2.7.21 [81,82]. The data are accessible through the NCI Genomic Data Commons (GDC) data portal (https://portal.gdc.cancer.gov). The GDC Data Portal provides access to the subset of TCGA data that have been harmonized (i.e., HTseq read mapping) against GRCh38 (hg38).

The aggregated data were pre-processed, normalized, and filtered prior to analysis, using different *TCGAbiolinks* functions. We pre-processed the data using the function *TCGAanalyze Preprocessing*, estimating the Spearman correlation coefficient among all samples. Samples with a correlation lower than 0.6 were identified as possible outliers and removed. It has been demonstrated that divergent tumor purity levels can lead to a false interpretation of differentially expressed genes between cancer and normal samples, as it may induce a confounding effect in the analysis of transcriptomic dataset [83]. To account for this possible effect, we filtered samples according to a derived consensus measurement of purity of 0.6 [83] as implemented in *TCGAbiolinks* [82]. We normalized the data to adjusts for external factors that were not of biological interest and to ensure that expression distributions of each sample were similar across the data. To this goal, we applied the function *TCGAanalyze Normalization*, implementing (i) within-lane normalization to adjust for GC-content effect on read counts [84] and (ii) between-lane normalization to adjust for distributional differences between lanes, i.e., sequencing depth [85]. Lastly, the data were full quantile filtered, using a threshold of 0.25, implemented in the function *TCGAanalyze Filtering* to remove features with low expression across the samples. We retained only samples containing the PAM50 intrinsic molecular subtypes, along with protein coding genes.

To explore the global structure of the high-dimensional dataset, we applied Principal Component Analysis (PCA) with the aim of (i) examining to what extent differential expression within the primary conditions of interest, could be distinguished, along with (ii) identifying possible batch effects. PCA was computed using the *prcomp* function from the R package *stats*. Exploratory analyses were undertaken on normalized log2 transformed read counts to relieve the heteroscedastic behavior of raw read counts. A pseudo-count of 1 was added to avoid taking the log of zero. Differential expression analysis was performed using the Bioconductor package *limma* [86]. *limma* integrates a range of statistical methods for effective analysis of gene expression experiments. At its core lies the ability to fit gene-wise (rows) linear models to the matrix of expression levels. This approach allows for flexibility in the sense that entire experiments as an integrated whole, can be analyzed, rather than step-by-step comparisons between pairs of treatments. Gene-wise linear models empower the sharing of information between samples, allowing one to model correlations that might be present between samples due to repeated measures or the presence of covariates. As of such, linear models allow for the adjustment of effects of multiple experimental factors or batch effects. The linear models describe how the coefficients (treatments) are assigned to different samples. Another important statistical component of *limma* is the empirical Bayes procedure, which facilitates the moderation of the gene-wise variances. This method estimates an optimal variance for each gene as a trade-off between the gene-wise variance, procured for that gene alone, and the global variance across all genes. *limma* linear modeling is conducted on log-CPM values, assumed to be approximately normally distributed and with an independent mean-variance relationship. It has been demonstrated that for RNA-Seq and other sequence count data, the variance is often dependent of the mean [87]. To remove heteroscedasticity, we applied the *voom* function, converting the mean-variance relationship through lowess fit and subsequently uses this to estimate gene-wise variances. For each gene, the inverse of the variance is then applied as “precision weight” in the downstream *limma* framework [49]. We adjusted for multiple testing using the Benjamini & Hochberg procedure of controlling the false discovery rate (FDR) or adjusted p-value. Significance was defined using an adjusted p-value cutoff of 0.05 together with a log-fold-change (logFC) threshold of 1 or −1 (for up- and down-regulated genes, respectively). Differentially expressed genes were visualized in a volcano plot, created using the *TCGAVisualize volcano* function of *TCGAbiolinks*. We included, directly in the design matrix for differential expression analysis, as a source of batch effect the information on the Tissue Source Site (TSS), upon exploration with PCA. We did not incorporate the effects of plate as we could see from the data, that the plates of interest were from the same TSS. Thus, the TSS was treated as a surrogate to avoid adding and extra parameter and associated degrees of freedom.

### Modeling of protein-peptide complexes

We modeled protein-peptide interactions with the scope of: (i) predicting the binding interface and the 3D structure of the complex between Bcl2a1 and Hrk BH3 regions, and (ii) identifying the location of the cancer mutations of Bcl2a1.

To model protein-peptide interactions, we applied comparative modeling, implemented in the program *MODELLER* v.9.15 [88], generating ten models for each alignment. *MODELLER* carries out comparative protein structure modeling by satisfying spatial protein structure restraints and optimizing the structure until a model that best satisfies the spatial restraints is acquired. In our modeling, we used as additional restraints the distance between V74 of the hydrophobic cleft of the template Bcl2a1 (chain A) and the invariant leucine for *h2* in each of the target BH3 peptides (chain B). To infer reliability and discriminate between models calculated from the same alignment, we applied statistically optimized atomic potentials, specially trained for scoring and assessing protein-peptide interaction [89]. We used the web server *VADAR* v.1.8 [90] to further assess the quality of the models. One model for each alignment was retained after these assessments. As a template structure, we used the known X-ray 3D structure of Bcl2a1 in complex with the BH3 peptide from the canonical BH3-only protein Puma (Bcl2-binding component 3, PDB ID 5UUL, R= 1.33 Å [60]). We generated the models of the complexes between Bcl2a1 and two BH3-like peptides or Hrk (Hrk_1, residues 28-50 and Hrk_2, residues 63-85).

### Identification of cancer mutations

We retrieved known missense mutations in the coding regions of BCL2-A1 and HRK using the *MuTect2* pipeline [91] for the TCGA-BRCA samples, which compares tumor to a pool of normal samples to find somatic variations. We used the pipeline as implemented in the *TCGAbiolinks* function *GDCquery_Maf*. We integrated this search with breast cancer mutations deposited in other studies available in *CBioPortal* [92] and *COSMIC* [93]. We also verified that the mutations which were not found in *ExAC* [94] as polymorphisms occurring in the health population.

### Structure-based prediction of the functional impact of mutations

We used the *FoldX* (http://foldxsuite.crg.eu) empirical force to predict changes in stability and interaction energies [95]. The *FoldX* energy function is obtained using a union of physical energy terms (e.g., van der Waals interactions, hydrogen bonding, electrostatics, and solvation), statistical energy terms, and structural descriptors that have been found important for protein stability. We used an in-house *Python* wrapper to support the systematic substitution of all wild-type residues to any of the 20 canonical amino acids, as recently applied to other cases of study [70,71,96,97]. With this tool we conducted *in silico* saturation mutagenesis, employing the *FoldX* energy function, predicting ΔΔ G values for all possible mutations in our modeled complexes. We applied the *RepairPDB* module from *FoldX*, optimizing the conformation of the model by repairing residues characterized by unfavorable torsion angles or, Van der Waals clashes. Subsequently, mutagenesis was carried out, applying the *BuildMode*l module from *FoldX*, independently mutating each residue at every position and calculating the Δ ΔG values. The prediction error of *FoldX* lies around 0.8 kcal/mol [95]. To infer the reliability of the predictions and discriminate between neutral and deleterious mutations, we applied a threshold of 1.6 kcal/mol (i.e., twice the prediction error) [97]. For visualization purposes, we truncated the ΔΔ G values a cutoff of 5 kcal/mol in the heatmaps. The cutoff was derived by investigating the distribution of experimental Δ ΔG values from the *ProTherm* database [98]. The vast majority of experimental ΔΔG values fall within −2.5 and 5 kcal/mol and as such *FoldX* predicted substitutions exceeding this value might be speculated to be overestimated. Other details on the saturation mutagenesis protocol are provided in our previous publication [97].

### Protein Structure Network Analysis

We used the *PyInteraph* suite [66] to derive a contact-based Protein Structure Network (PSN) [67] for the complex of BCL2-A1 with BH3 peptides. We used as a reference starting structure the structure of the complex of BCl2-A1 with PUMA, which has been solved by X-ray. Since the *PyInteraph* method has been designed to work on a structural ensemble, we collected a representative ensemble of ten conformations for the complex using *CABS_Flex 2.0* [69].

We considered as interacting pairs for the PSN any two residues whose side-chain centers of mass lied within 5.0 Å. This cut-off was selected as suggested by a recent benchmarking of the method [67] We also applied a 20% cutoff to the persistence (pcrit) of the interaction in our PSN to filter out transient and spurious interactions from our network, as previously suggested [66,99]. We included all the residues with the exception of glycine for the contact analysis. We applied a variant of the depth-first search algorithm to identify the shortest paths of communication, whereas hubs were defined as residues with a degree of more than 3 (i.e., linked by more than three edges in the network), as generally applied to PSNs [100].

## Acknowledgements

The results shown here are in part based upon data generated by the TCGA Research Network: https://www.cancer.gov/tcga. The calculations described in this paper were performed using the DeiC National Life Science Supercomputer Computerome at DTU (Denmark).

