## Supplementary figures and images for "Alterations of the pro-survival Bcl-2 protein interactome in breast cancer at the transcriptional, mutational and structural level"

### Figure S1

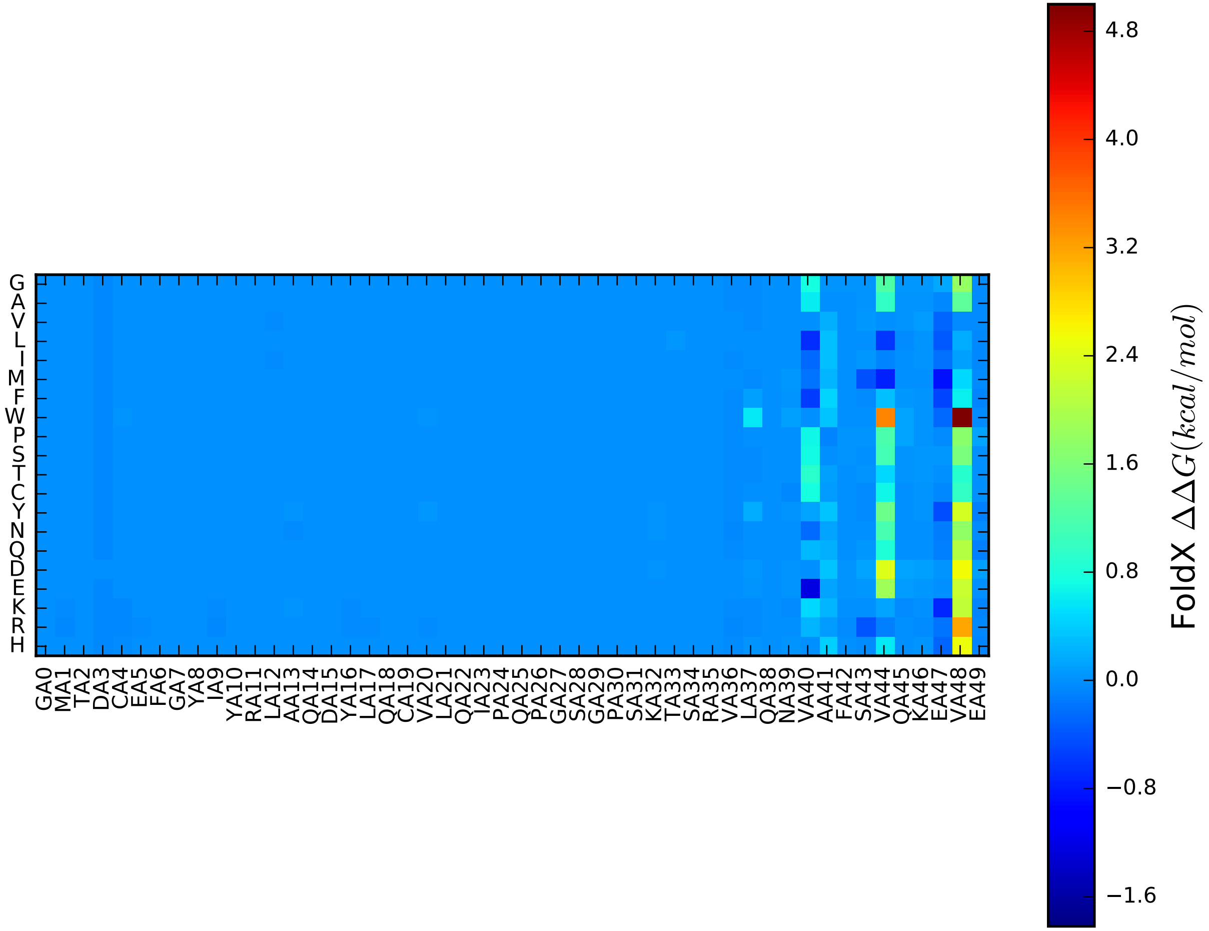

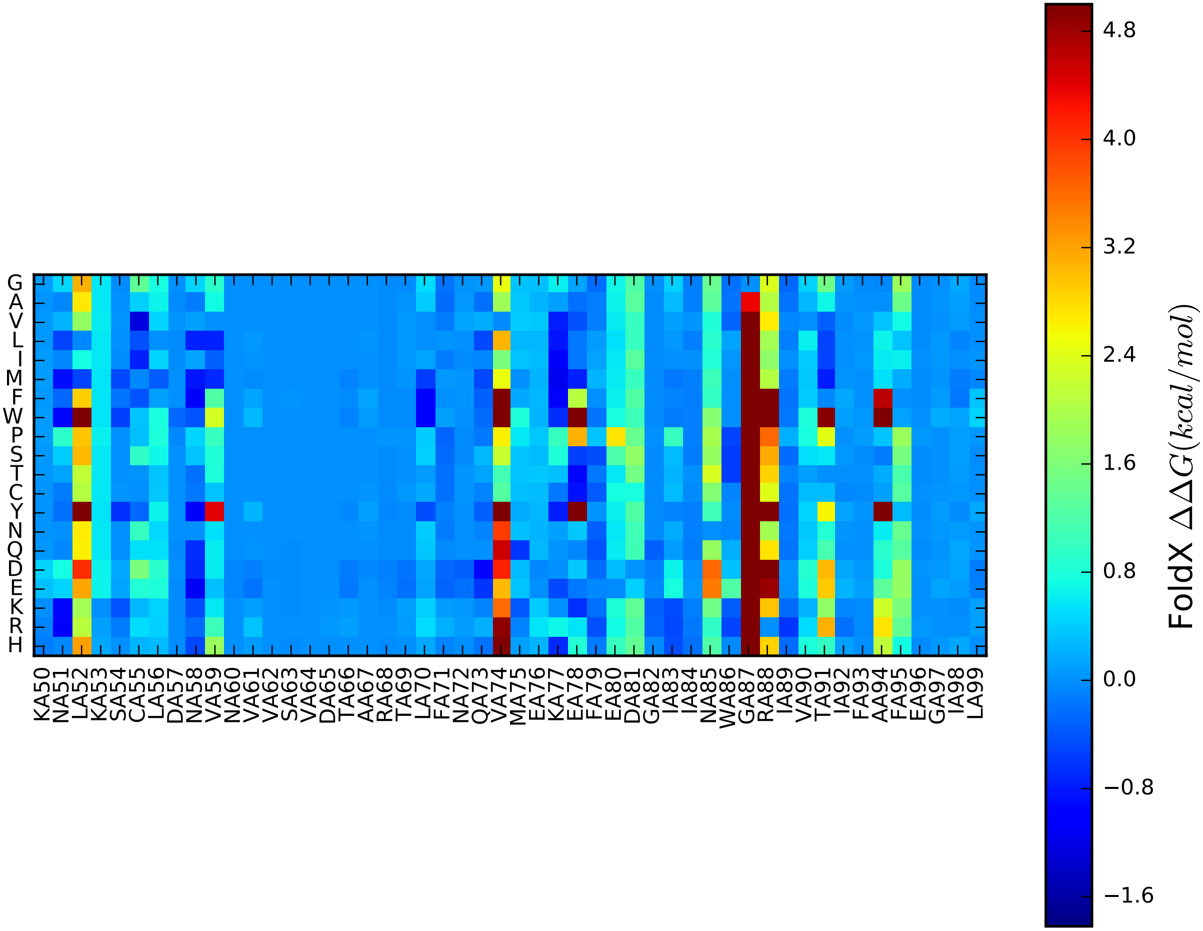

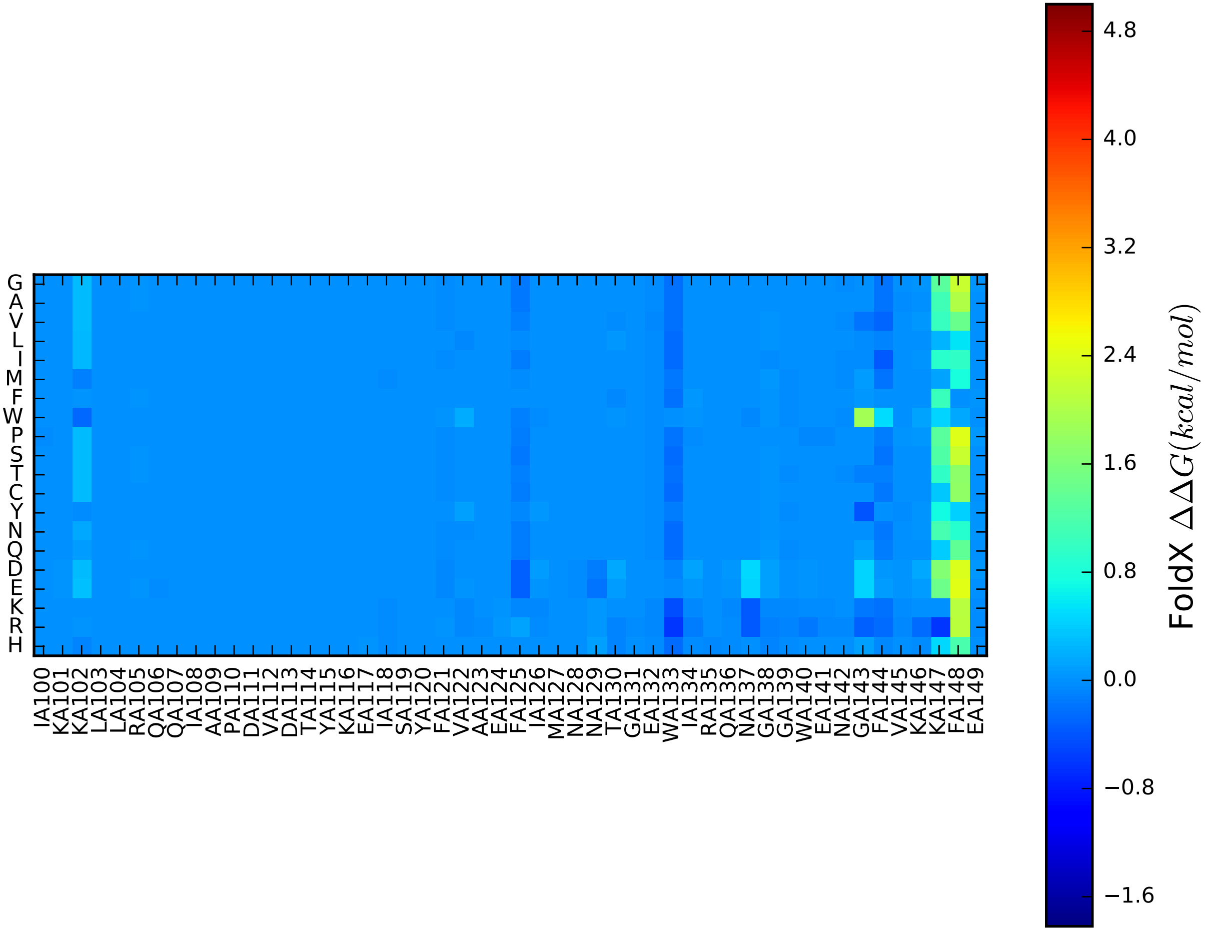

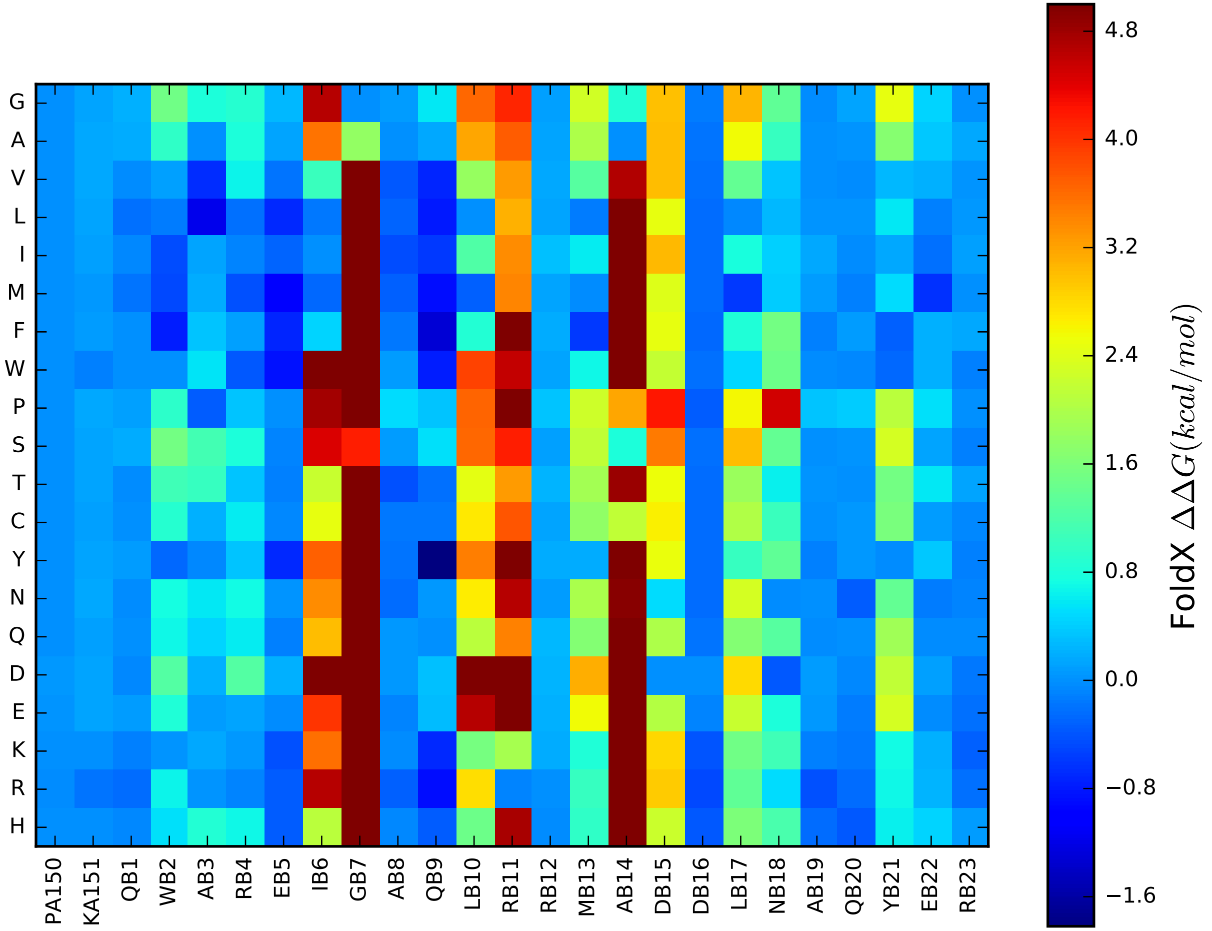
